# Self-organized coexistence of phage and a population of host colonies

**DOI:** 10.1101/2024.09.02.610744

**Authors:** Anjali Yadav, Namiko Mitarai, Kim Sneppen

## Abstract

Phages and bacteria coexist under widely different conditions, ranging from liquid cultures to oceans, soil, and the human gut. However, our models are typically limited to well-mixed liquid cultures governed by mass-action kinetics. Here, we suggest a modification to the Lotka-Volterra dynamics by including the formation of microcolonies. By analyzing the model in an open system with a steady influx of bacteria, we predict that the colony size distribution is power-low distributed with steeper exponents for the stronger external influx. In the realistic case where the phage attack rate to individual colonies is proportional to their radius, we obtain self-organization to a steady state where the maximal colony size is smaller for stronger external driving.

---

The coexistence of bacteria and its virulent phage was predicted already by Campbell [1] and later confirmed experimentally in continuous chemostat culture [2, 3] as well as by serial transfer [4]. However, the predator-prey relationship between phage and host is destabilized by the many off-spring phages from each infected bacterium, suggesting that well-mixed conditions should have extreme oscillations that may easily lead to the extinction of the host population [5]. These extreme oscillations are often not observed in experimental data [2], perhaps because the bacterial liquid culture tends to develop heterogeneity associated with some bacteria growing on the glass walls of the chemostat [4]. Here, we will introduce a model for another spatial heterogeneity, using the fact that many bacteria types tend to grow in clusters and that these clusters provide some spatial refuge against phages [5–7].

As a wide-ranging goal, we aim to develop a better understanding of diversity and coexistence within ecological systems. Commonly, researchers employ mean-field models to interpret the associated dynamics, using generalizations and extensions of the famous Lotka-Volterra (LV) framework [8, 9]. While these equations have given us much insight into concepts like the competitive exclusion principle [10–14], they tend to be too fragile to support the coexistence of many competing species. To remedy these deficiencies, Robert Holt [15] suggested that spatial heterogeneity is important for stabilizing ecosystems. From this perspective, the current paper explores conditions where clusters of prey may self-organize to help their population evade predators more effectively, thereby stabilizing their overall numbers.

We initiate our study by considering the standard LV form of predator-prey modelling, which mathematically studies the temporal evolution of bacterial *B*(*t*) and phage *P* (*t*) populations in a well-mixed system. The LV equations implicitly assume that each predator is equally likely to encounter any prey individual. We extend the conventional LV framework by suggesting a model where each bacterium can develop into a distinct microcolony as bacteria grow, thereby introducing a microcolony-based modification to the interaction dynamics. This represents a shift from a model of distributed individual bacteria and phages to one featuring distributed microcolonies and phages.

## Model

A schematic representation of the model is illustrated in Fig. 1, where each blue sphere symbolizes a microcolony of bacteria. We use a well-mixed model of microcolonies and phages in the system, disregarding other spatial considerations. The system has an influx of individual bacteria with a rate *α* per unit volume. Each bacteria being added to the system may subsequently grow into a microcolony. Bacteria in microcolonies have a constant growth rate of *λ*, implicitly assuming a constant resource per bacteria. The rate at which a phage infects a colony depends on the size of the colony (*B*), infection rate (*η*) and phage density (*P*). We assume the rate is given by *ηPB*^*ν*^, where we later motivate the value for the exponent *ν*. Each infection event results in an instantaneous colony burst with the release of *β* (burst size) phages per bacterium of the colony. This is a simplification where we ignore the latency time for the phage production and assume that the phages produced by infected bacteria immediately reinfect other bacteria in the system to burst all the cells eventually. Finally, the free phage decays slowly at a constant rate *δ*. Thus, the system is slowly driven by bacteria addition and dissipated by sudden colony infection and collapse.

**FIG 1.**
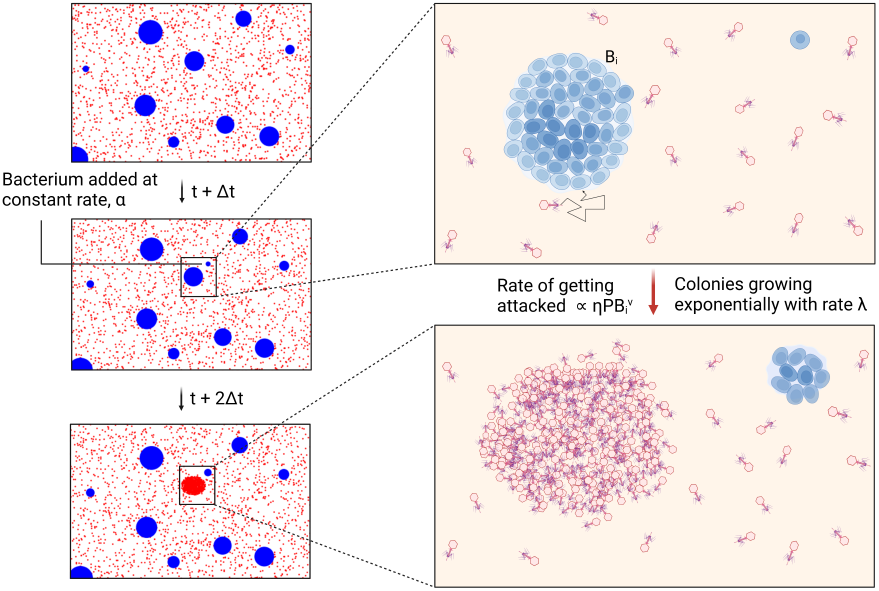
Schematic description of the model. A new bacterium is introduced to the system at a rate *α* per volume, and each bacterium grows exponentially at a rate *λ* to form a colony (blue spheres). A colony of population *B*_*i*_ is attacked by a phage (red dots, density *P*) at a rate 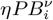. The phage attack results in all the bacteria in the colony eventually being infected, producing *βB*_*i*_ new free phages. Free phage decays at a rate *δ*.

We simulate the dynamics of phage-bacteria interactions in a system where each bacterial colony *i* (*i* = 1, …, *N*) is modelled as a discrete entity. The overall phage density *P* and the population size of bacteria within each colony *B*_*i*_ are treated as continuous variables, and phage decay and colony growth is simulated deterministically. The simulation proceeds with a discrete time step Δ*t*. During each time step, in addition to the deterministic growth of each colony *B*_*i*_(*t* + Δ*t*) = *B*_*i*_(*t*)*e*^*λ*Δ*t*^ and phage decay *P* (*t* + Δ*t*) = *P* (*t*)*e*^*−δ*Δ*t*^, the following stochastic event can happen: 1) A single bacterium is added to the system with probability *α*Δ*t* per unit volume, starting a new colony and 2) Each colony *i* is subject to infection events with probability 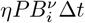. When the infection happens, the colony is removed, and *βB*_*i*_ phages are produced The algorithm is presented in the Supplementary Material (SM Sec. S1).

We study the system for two values of *ν*. The simple *ν* = 0 represents the colony infection rate independent of colony size. However, in a well-mixed liquid culture, a colony’s attack rate is expected to be determined by diffusion-limited search. If so, the adsorption rate is proportional to the radius of the target, which for a colony of size *B* would scale as *B*^1*/*3^ [16]. It was also confirmed experimentally [17]. Hence, we consider the *ν* = 1*/*3 case. We discuss other values of *ν* briefly at the end.

Unless otherwise stated, we use *η* = 10^*−*8^*ml/h, λ* = 2*h*^*−*1^, *δ* = 0.1*h*^*−*1^, *β* = 100. These values are realistic ranges for typical *E. coli* phages [18, 19]. The time step is set to Δ*t* = 10^*−*4^ h, which is small enough to ensure that the probability for stochastic events per Δ*t* does not exceed 1. The stochastic simulation is done within a total volume *V*, where *V* = 1*L* unless otherwise noted. The value of influx *α* is varied in the simulations.

### Population dynamics

Model dynamics are shown in Fig 2. For comparison, panel (a) illustrates characteristic oscillations of the LV model (SM eq. (S1)) where bacteria do not form colonies. Panel (b) shows the dynamics when considering *ν* = 0, where episodic population dynamics are associated with the collapse of individual large colonies. Panel (c) presents the more realistic case with *ν* = 1*/*3. Here, the populations are stabilized provided *V* is large enough.

**FIG 2.**
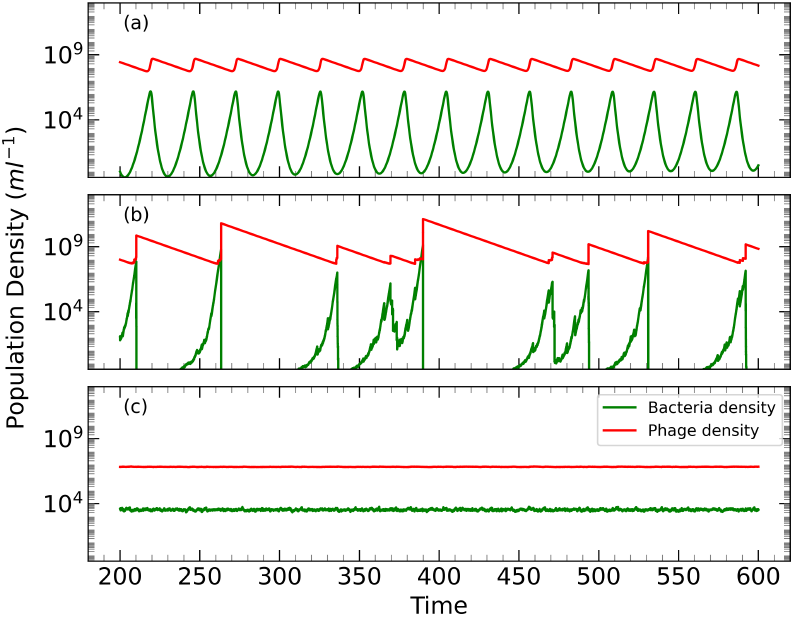
Temporal dynamics of bacterial and phage population density simulated using the Gillespie algorithm for three different models: (a) the Lotka-Volterra (LV) model, (b) the model with *ν* = 0, and (c) the model with *ν* = 1*/*3, illustrating more stable population densities. Phage parameters are *η* = 10^*−*8^, *λ* = 2*h*^*−*1^, *δ* = 0.1*h*^*−*1^, *β* = 100. Simulations use *V* = 1*L* and *α* = 0.1*h*^*−*1^*ml*^*−*1^. The near-constant dynamics in panel C) are replaced by episodic bust-boom dynamics for smaller bacterial influx *α · V ∼* 1*/h* (See Fig. S5).

Panel b) shows a repeated pattern of bacterial population growth followed by sudden drops associated with phage attacks. Complementary to these, one observes sudden increases in phage density followed by exponential decay set by *δ*. When the phages eventually infect a sufficiently large colony, there is a sudden spike in phage density, which hugely increases the attack rate on all other colonies, leading to their nearly simultaneous collapse. Subsequently, the many phages prevent even the smallest colonies from surviving. Once the phage density decreases to a sufficiently low level, the colonies can begin to grow again, and the cycle repeats.

The dynamics stabilize for *ν* = 1*/*3, as seen in panel c). This is because larger colonies are more easily found and eliminated by the phages; they grow but are eliminated before they become substantial. In other words, the phages keep the system in check and prevent any single colony from dominating, preventing the extreme spike of the phage density and reducing the fluctuation. At the same time, the stochastic colony dynamics prevent the LV oscillation. Notably, the stochastic colony dynamics becomes visible when the number of colonies in the system is small. This happens obviously for a small enough volume *V*, and in addition expected for small values of influx *α* or phage adsorption rate *η* (SM Fig. S10).

An important model parameter is the rate of addition of individual bacteria *α*. Figure 3 shows how *α* influence the densities of phages (*P*), bacteria (*B*), and number of colonies *ρ* per unit volume for the *ν* = 1*/*3 case. Also, the figure shows the average size ⟨*s*⟩ of the colonies. Interestingly, *P* and *B* varies less with *α* than ⟨*s*⟩ and *ρ*. As *α* increases, then ⟨*s*⟩ decreases, and *ρ* increases, balancing the overall population. Thus, the system self-organizes to maintain steady-state populations which are remarkably robust to variation in *α*.

**FIG 3.**
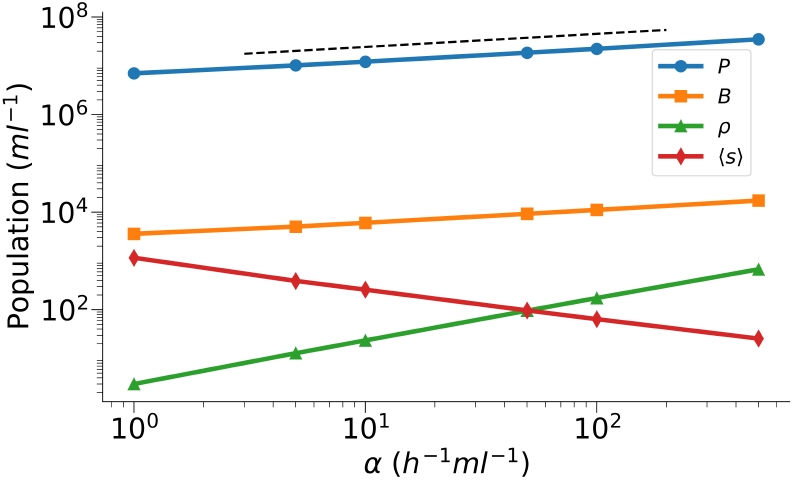
Dependence of average phage population density (*P*), average bacterial population density (*B*), average colony density *ρ*, and average colony size ⟨*s*⟩ on the bacteria introduction rate *α* for a system of *V* = 10 ml for *ν* = 1*/*3. The graph illustrates the inverse relationship between *α* and both bacterial population and density while showing an increase in average colony size with higher *α*, highlighting the complex dynamics of bacterial-phage interactions. The dashed line shows the analytical prediction of *P* dependence on *α* in small *α* limit, *P ∝ α*^1*/*4^.

### Colony size distribution

In infinite system limit, the density of colonies of size *s* at time *t, n*(*s, t*) can be described by

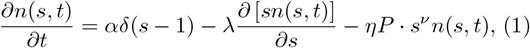

where *δ*(*x*) is the Dirac’s delta function. The phage density *P* then obeys

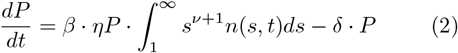

The *ν* = 0 model does have a steady state phage density. However, at any particular instant, *t* simulations show a probability distribution of the colony size

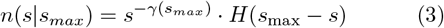

where *H*(*x*) is the Heaviside step function and *s*_*max*_ varies with time *t*. The dependence *γ*(*s*_*max*_) comes from large *s*_*max*_ being associated with periods that have small *P* and surviving colonies from increasingly separated introduction times. The decline of *γ*(*s*_*max*_) with *s*_*max*_ is explored in SM Fig. S1c. The time-averaged *n*(*s*) is

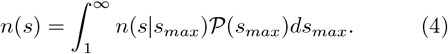

where 𝒫(*s*_*max*_) is probability distribution of *s*_*max*_. We evaluated 𝒫(*s*_*max*_) from the simulation data (SM Fig. S1c) as 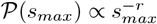 and integrated eq. (4) (SM Sec. S2.A). This gave us a power-law distribution

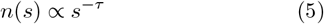

with a logarithmic correction. The simulation data shows power-law distributions over 4 decades with exponent *τ* between 1.3 and 1.5 depending on *α* (Fig. 4a).

**FIG 4.**
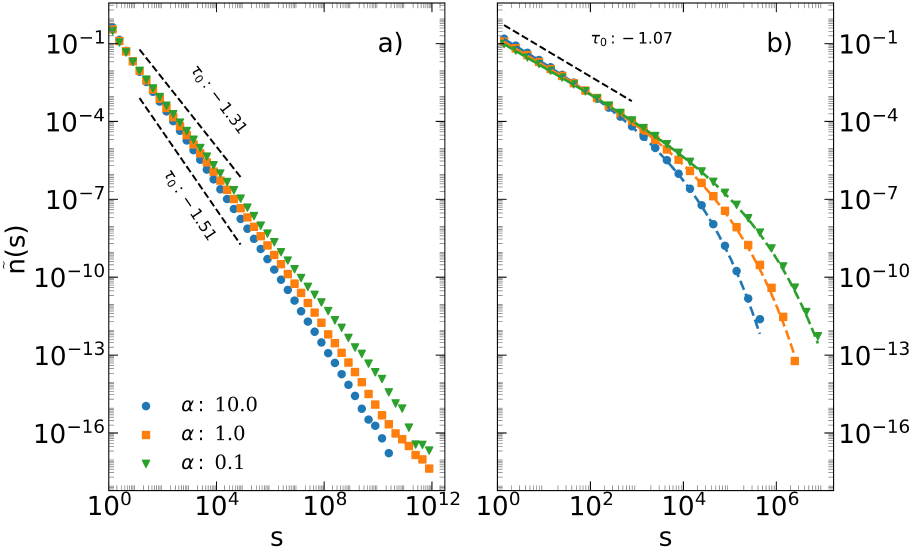
Normalized time-averaged colony size distributions ñ(*s*) = *n*(*s*)*/ ∫ n*(*s*)*ds* in the steady state for different values of the bacteria influx rate *α*. (a) Model with *ν* = 0. (b) Model with *ν* = 1*/*3. Coloured dashed lines in (b) represent the analytical solution eq. (7) for different *α* values.

For *ν* = 1*/*3, the system reaches a steady-state. Ignoring the fluctuations in *P* and solving equation eq. (1) for a steady-state for *s >* 1, we obtain

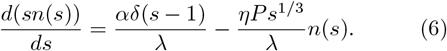

Solving this gives us (SM Sec. S2.B)

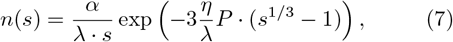

and the steady state condition for eq. (2) determines the value of *P*. The equation indicates a power-law decay, *n*(*s*) *∝ s*^*−*1^ moderated by the size-dependent cut-off reflecting increased vulnerability to phage attacks as colony size increases.

The analytical result eq. (7) shows an excellent agreement with numerical simulations as shown in Fig. 4b, when the value of *P* is numerical evaluated from the analytical expression in SM eq. (S14). For small *α* limit eq. (S14) predicted *P ∝ α*^1*/*4^, which also agrees with the result in Fig. 3.

## Discussion

Inspired by phage-bacteria systems, this paper explored LV dynamics with the prey population partitioned into colonies. When bacteria grow in micro-colonies, the phage has more difficulty in locating them, though once they succeed, the phage predates the entire local colony [20, 21]. If all the colonies were the same size, this would reduce to an ordinary LV dynamics of colonies and phages with larger adsorption and larger burst sizes (eq. S2), resulting in oscillating population sizes. However, in our model, variation between colony fates disrupts synchronous oscillation. If the chance for a phage to find a colony was independent of colony size, the model predicts a power-law distribution of colony size *s, p*(*s*) *∝* 1*/s*^1.5^. With a more realistic diffusion-limited search, the rate to locate the colony grows with *s*^1*/*3^ [17, 22]. This results in a finite cut-off of the colony sizes (Fig. 4), which in turn reduces the episodic fluctuations towards zero as system size increases (Fig. 2).

The modelled system self-organizes the sizes and number of colonies to total populations that only change weakly with bacteria influx *α* (Fig. 3). Further, the value of the phage-to-bacterial ratio is independent of *α*. Importantly, we also considered a model version where bacteria cells disperse stochastically from existing colonies’ surface to start a new colony (Supplemental Material Sec. S3). This realistic variation behaved similarly to our standard model, with the dispersion rate from existing colonies replacing the influx rate *α*. This underscores the robustness of the obtained self-organization.

The *ν* = 1*/*3 scaling assumes a colony to be densely packed, while the simplest *ν* = 0 case could realized if cells formed a loosely dispersed aggregate of a roughly constant volume. However, bacterial aggregate can have various geometry [21]. Therefore, we also considered different *ν* values in Supplemental Material, Sec. S4. For fixed influx rate *α* we find that both *B* and *P* decrease with increasing *ν* while *P/B* remain constant. This reflects that larger *ν* increased the attack rates on colonies. For *ν* = 1, the bacteria only grow to rather small colonies (a hundredfold lower) while still providing heterogeneity that suppresses oscillations (Fig. S4a).

Noticeably, the obtained *P/B* ratio from Fig. 3 is close to the LV prediction of *λβ/δ* = 2000. This value is larger than the classically reported *P/B* ratio of order 10 [23]. This *P/B* ratio is also obtained in the mean-field version of our model as the colonies essentially act as a changed *η* which rescale *P* and *B* equally, see Sec. S1 of Supplemental material. A smaller *P/B* ratio can be expected by smaller *λ, β*, and/or larger death rate *δ*. In the nutrient-poor ocean water, *λ* and *β* are smaller than used here, while our used *δ* is close to the one reported by [19].

In addition, there are idealizations in our colony model which systematically overestimates the obtained *P/B* ratio.

First, we neglected that phages escaping from one colony must travel some distance to locate a new colony. This gives more time for colonies to grow, while phages can decay while travelling. We simulated this situation by considering spatiotemporal dynamics in two dimensions with phages diffusing between fixed colonies [24] (SM Sec. S5). For a small *α* where the colony density is low, the predicted ratio of *P/B* easily becomes lower than 10.

Second, our model neglected that phages spreading in a colony result in a time delay of phage release. A minimal time delay may be estimated by simulating a version of the model where the virus spreading within a colony is assumed to be well mixed, resulting in an exponential decay of colony size over time (SM Sec. S6). This reduces the severity of the phage attack. Conservatively assuming the colony collapse rate to be ln 10/h suggests a factor 2 decrease in the *P/B* ratio (SM Fig. *S*6, *S*7).

Finally, the spread of infection is further limited by superinfected cells close to the colony surface [7, 25, 26] and/or hindered by an extracellular matrix which can trap phages [27]. Such limits on the phage predation naturally reduce *P* and increase *B*, again bringing the *P/B* ratio down. In simplest approximation, at least by a factor of two.

In conclusion, phages predating bacteria in colonies should result in lower *P/B* ratios than phages predating freely living bacteria.

The feature of hiding the prey population in clusters has analogues in other ecosystems. Fish aggregate in schools to minimize predator attacks [28], while cicadas satiate their predators during rare but synchronous swarming events [29]. Our simplest scenario considers an extreme version of heterogeneity, where the predator’s ability to localize a colony was independent of its size. This perspective of replacing an attack rate dependent of *size*^1*/*3^ with a constant is analogue to the difference between the “dynamic kill the winner” model of [30] and the simple Boom-Bust model of [31]. Taking into account the phage versus colony dynamics, our model enhances the magnitude and duration of the “busts” because the collapse of a large colony is followed by a prolonged period of phage dominance with the associated elimination of bacteria.

Considering the large diversity of coexisting phages and their host observed in the ocean [32], our model suggests this is to be driven by individual bacteria species that boom as colonies grow on a localized nutrient, followed by a bust due to a rare infection by a scarce phage. Ref. [32] showed that, for the *Vibrio* species and their phages, the snapshots of the populations of individual phage species are often less than 0.1 phages per ml, but sometimes count up to a few hundreds of phages per ml. Given each *Vibrio* species was on average *<* 1 cells per ml [33], this is consistent with the large stochasticity in our model when the number of colonies in a sample is limited. It is also worth mentioning that colony or aggregate formation and dispersal are often quorum sensing regulated [34, 35], allowing the bacteria to regulate the colony size depending on the environmental clues.

In summary, we introduced a minimal stochastic model for phages spreading between host colonies. The model highlights the self-organization of colony sizes and suggests further studies of both phage propagating between colonies and phage spreading within a colony. In the case of colony-forming bacteria in the ocean, this include hydrodynamic mixing above the Kolmogorov scale, while spreading within a colony depends on the phage’s ability to propagate in dense environments. We hope this work will stimulate further research on the interplay between phages and bacteria that form or leave dense environments.

## Acknowledgment

This research was funded by the Novo Nordisk Foundation (NNF21OC0068775) and the Danish National Research Foundation (grant no. DNRF170).

