## Supplementary text for "Self-organized coexistence of phage and a population of host colonies"

**S7. Fluctuations in $B$, $P$ and colony sizes with adsorption rate, $\eta$ and influx rate, $\alpha$** . . **11**

### S1. LV model, Algorithm, and the mean-field models

#### A. Modified Lotka-Volterra Model for Bacteria and Phage Dynamics

The bacterial ($B$) and phage ($P$) populations are modeled using a modified Lotka-Volterra framework:

$$\begin{aligned} \frac{dB}{dt} &= \alpha + \lambda B - \eta BP, \\ \frac{dP}{dt} &= \beta\eta BP - \delta P. \end{aligned} \tag{S1}$$

Here, $\alpha$ is the bacterial influx rate, $\lambda$ is the bacterial growth rate, $\eta$ is the infection rate, $\beta$ is the burst size per lysis event, and $\delta$ is the phage decay rate.

#### B. Stochastic Simulation Algorithm

We simulate the dynamics of phage-bacteria interactions in a system where each bacterial colony $i$ ($i = 1, \ldots, N$) is modelled as a discrete entity. The phage-bacteria system evolves in discrete time steps $\Delta t$, where $\Delta t \ll 1/\max(\text{rate})$ ensures numerical stability. The simulation at each time step proceeds as follows:

1. **Update Bacterial Growth:** Each bacterial colony $i$ ($i = 1, \ldots, N$) grows deterministically:

$$B_i \to B_i \exp(\lambda \Delta t),$$

2. **Update Phage Decay:** The overall phage density $P$ decreases deterministically:

$$P \to P \exp(-\delta \Delta t),$$

3. **Simulate Stochastic Colonization:** With probability $\alpha \Delta t$ per unit volume $V$, a new bacterial colony is introduced into the system. If colonization occurs, a colony of one bacterium is added to the system.

4. **Simulate Phage Infection:** For each bacterial colony $i$, calculate the probability of infection:

$$\text{Infection Probability} = \eta P B_i^{\nu} \Delta t,$$

   where $B_i^{\nu}$ represents a nonlinear infection dependence on colony size.

   - If an infection occurs, colony $i$ is lysed ($B_i \to 0$), and $\beta B_i$ new phages are released into the system.

5. **Repeat Steps 1–4:** Repeat these steps at each time step until the system reaches the total simulation time.

#### C. Modified LV Model with Fixed Colony Size $s_0$

The model considers a system of colonies, each with a fixed size $s_0$. Colonies grow and divide into two at a rate $\lambda$. The infection rate is assumed to depend on the radius of the colony, which scales as $s_0^{1/3}$, as outlined in the main text. An influx term, $\alpha_s$, is also introduced, which is proportional to $\alpha/s$.

Let $n$ denote the number of colonies of size $s_0$ in the system. Under these assumptions, the modified Lotka-Volterra equations are expressed as follows:

$$\begin{aligned} \frac{dn}{dt} &= \alpha_s + \lambda n - \eta {s_0}^{1/3} n P, \\ \frac{dP}{dt} &= \beta s_0 \ \eta {s_0}^{1/3} n P - \delta P. \end{aligned} \tag{S2}$$

##### D. Mean field model and $P/B$ ratio in the steady state.

Suppose the density of colony size $s$ is $n(s,t)$. Since the chance of a bacterium belonging to a size $s$ colony is $sn(s,t)$, the mean-field approximation for the total bacteria density ($B = \int_1^\infty sn(s,t)$) and the phage density $P$ with $\alpha = 0$ gives

$$\frac{dB}{dt} = \lambda B - \eta P \int_1^\infty s^{\nu+1} n(s,t) ds, \tag{S3}$$

$$\frac{dP}{dt} = \beta\eta P \int_1^\infty s^{\nu+1} n(s,t) ds - \delta P. \tag{S4}$$

In the steady state we obtain $P/B = \lambda\beta/\delta$, which is the same expectation of the steady state for the ordinary LV model eq. (S1) with $\alpha = 0$.

### S2. Analytic Framework for Bacterial Colony Size Distribution

#### A. The evaluation of the $n(s)$ distribution for $\nu = 0$ case

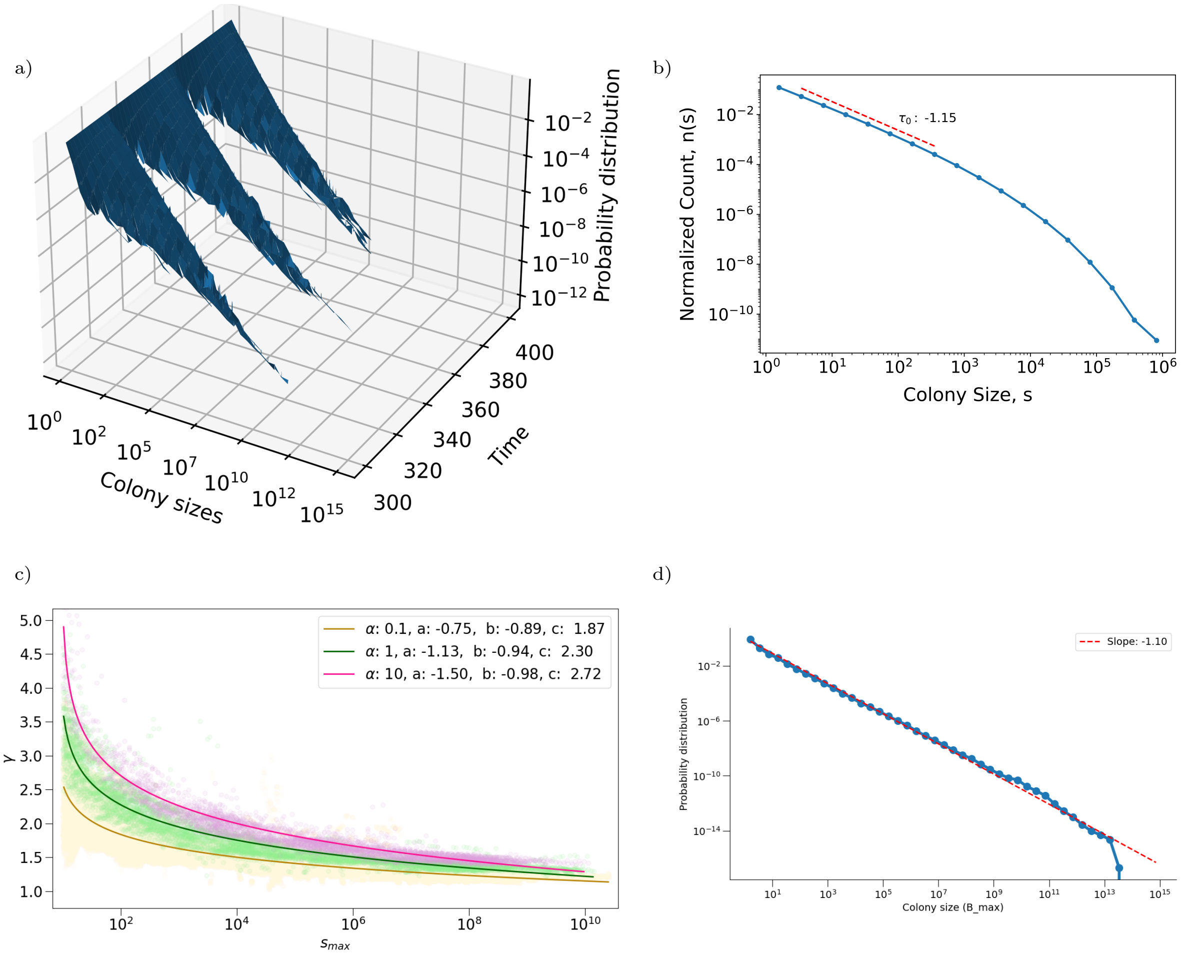

Figure S1: a) Probability distribution of colony size over time for $\alpha = 1$, $V = 1\,\mathrm{L}$. b) Probability distribution of maximum colony sizes at steady state for $\alpha = 1$, $V = 1\,\mathrm{L}$, which fits well to $s_{max}^{-r}$ with $r \sim 1.1$ for the range of $\alpha$ explored. c) $\gamma$ as a function of $s_{max}$ for different values of $\alpha$. The lines show the fit to the function $\gamma = a\log(\log(s_{\max}) + b) + c$ d) Probability distribution of maximum colony sizes ($s_{max}$) at steady state, which fits well with $r \sim 1.1$ for the range of $\alpha$ we are exploring.

We found that the $s_{max}$ dependence of $\gamma$ is fitted well by the function $\gamma = a\log(\log(s_{\max}) + b) + c$. Starting from the integral in the main text,

$$n(s) = \int_1^\infty s^{-a\log(\log(s_{\max})+b)-c} H(s_{\max} - s) s_{\max}^{-r}\, ds_{\max} \tag{S5}$$

we substitute $s_{\max} = 10^n$ and $s = 10^m$. This substitution simplifies the integral to:

$$n(s) \propto s^{-c} \int_{m+b}^\infty n^{-ma} e^{(1-r)\ln(10)n}\, dn$$

Further simplification leads to:

$$n(s) \propto s^{-c} s^{a \log_{10}((r-1)\ln(10))} \int_{f(m,b)}^{\infty} t^{-ma} e^{-t}\, dt$$

where $f(m,b) = (r-1)\ln(10)(m+b)$. The integral can be expressed in terms of the incomplete gamma function and substituting $r \sim 1.1$ (see Fig.S1d):

$$n(s) \propto s^{-c-0.64a} \int_{f(m,b)}^{\infty} t^{-ma} e^{-t}\, dt$$

Thus, we obtain:

$$n(s) \propto s^{-c-0.64a} \Gamma(-ma+1, f(m,b)) \tag{S6}$$

For $(-ma+1) > 0$, the incomplete gamma function $\Gamma(-ma+1, f(m,b))$ can be expanded as:

$$\Gamma(-ma+1, f(m,b)) = (-ma)! e^{-f(m,b)} \sum_{k=0}^{n} \frac{f(m,b)^k}{k!} \tag{S7}$$

Since the expansion contains terms of order $\log(s)$, the primary contribution to $n(s)$ comes from,

$$n(s) \propto s^{-c-0.64a-(r-1)}. \tag{S8}$$

#### B. Analytical solution of the colony size distribution and Phage density for $\nu = 1/3$ case

Solving Eq. (6) in the limit $\alpha \to 0$, we obtain:

$$n(s) = \frac{\mathcal{N}}{s} \exp\left(-\frac{3\eta P}{\lambda} s^{1/3}\right), \tag{S9}$$

where $\mathcal{N}$ is the normalization constant. We use the variation of the coefficient method by assuming $\mathcal{N}$ can depend on $s$. Substituting eq. (S9) into $sn(s)$, the derivative is:

$$\frac{d(sn(s))}{ds} = \frac{d\mathcal{N}}{ds} \exp\left(-\frac{3\eta P}{\lambda} s^{1/3}\right) - n(s) \frac{\eta P}{\lambda} s^{1/3}. \tag{S10}$$

Inserting this with Eq. (6) gives

$$\frac{d\mathcal{N}}{ds} = \frac{\alpha\delta(s-1)}{\lambda} \exp\left(\frac{3\eta P}{\lambda} s^{1/3}\right). \tag{S11}$$

Integrating this from 1 to $s > 1$ and using $\mathcal{N} = 0$ when $\alpha = 0$ in the steady state, we obtain

$$\mathcal{N} = \frac{\alpha}{\lambda} \exp\left(-\frac{3\eta P}{\lambda}\right). \tag{S12}$$

Thus, the steady-state distribution $n(s)$ is fully determined by substituting Eq. (S12) into Eq. (S9). Assuming fluctuations in phage density are negligible, the value of $P$ is determined by the steady-state condition. We start with equation (2):

$$\beta\eta \int_{1}^{\infty} s^{\nu+1} n(s)\, ds = \delta.$$

Substituting the expression for $n(s)$ for $\nu = 1/3$ and solving the integral using integration by parts, we obtain:

$$\frac{3\alpha\beta\eta}{\lambda\delta a_0^4} \left(a_0^3 + 3a_0^2 + 6a_0 + 6\right) = 1, \tag{S13}$$

where $a_0 = \dfrac{3\eta P}{\lambda}$.

Rewriting the left-hand side (LHS) of this equation as a function of $P$, the following expression is obtained:

$$f(P) = \frac{3\alpha\beta\eta}{\lambda\delta}\left[\frac{\lambda}{3\eta P} + 3\left(\frac{\lambda}{3\eta P}\right)^2 + 6\left(\frac{\lambda}{3\eta P}\right)^3 + 6\left(\frac{\lambda}{3\eta P}\right)^4\right]. \tag{S14}$$

This relation is shown in Fig. S7b, where $f(P)$ is plotted as a function of $P$ to analyze the mean phage density of the system under varying influx rates.

Since $P$ decreases with $\alpha$, for a small enough $\alpha$ the last term of eq. (S14) becomes the most dominant, predicting $P \propto \alpha^{1/4}$ in the limit.

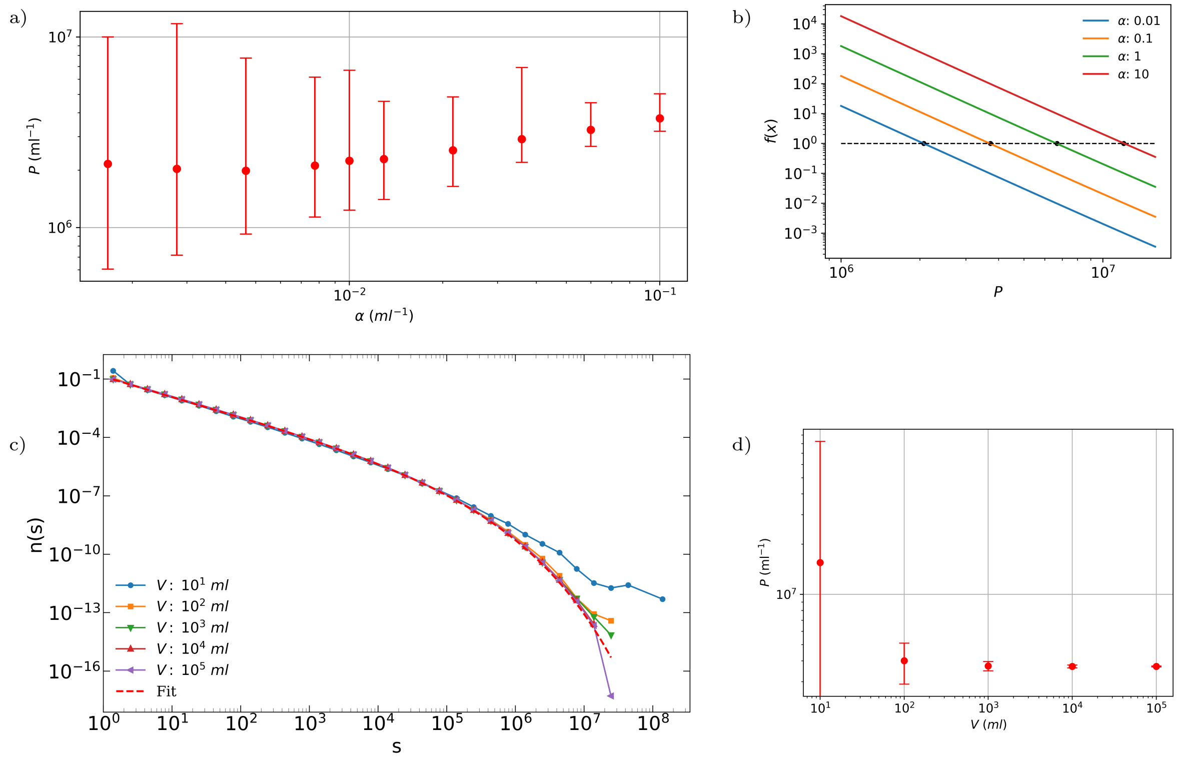

Figure S2: a) Mean phage density, $P$ and fluctuations in phage density as a function of the influx rate ($\alpha$), for a fixed system volume ($V = 1L$), all other system parameters are the same. b) Analytical calculation of the mean phage density using the steady-state equation obtained above. The analytically obtained value closely approximates the phage density obtained from numerical simulations. c) The colony size distribution for varying system volumes. As concluded analytically, the distribution is governed by the mean phage density. d) the mean phage density remains approximately constant for larger system volumes, leading to similar colony size distributions across these volumes.

#### S3. Model with bacterial dispersal from colonies

To model a more realistic system, the influx of bacteria is replaced by the dispersal of bacteria from existing colonies. In this scenario, each bacterium at the surface of a colony has a probability of leaving the colony, governed by a dispersion rate ($\gamma$). Dispersed bacteria grow into new colonies upon leaving their original host colony, thereby driving the emergence of new bacterial populations within the system. An additional step is included in the previous model to account for bacterial dispersal from existing colonies. Each bacterial colony $i$ has a probability of dispersing bacteria during a time step, given by:

$$\text{Dispersion Probability} = \gamma B_i^{2/3}\Delta t,$$

where $\gamma$ represents the dispersion rate. When a bacterium disperses from the $i^{\text{th}}$ colony, the colony population is updated as:

$$B_i \to B_i - 1.$$

Dispersed bacteria are considered to form new colonies, analogous to the bacterial influx ($\alpha$) in the original model.

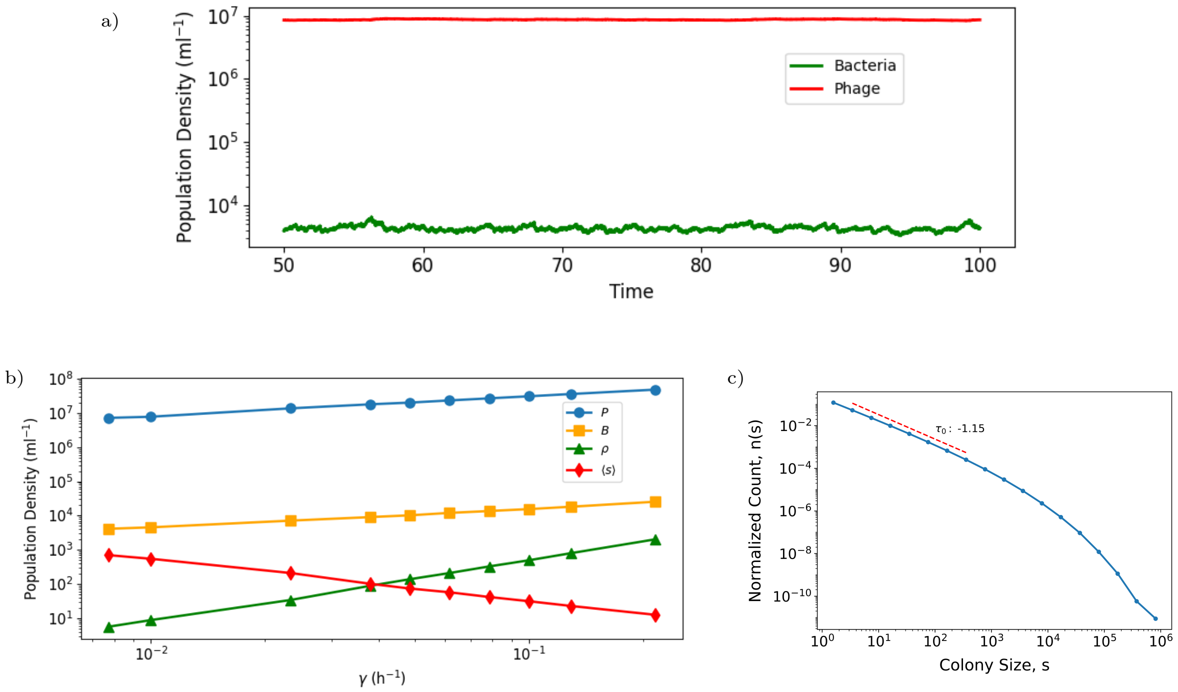

Figure S3: a) The steady-state dynamics of phage and bacterial populations for $\gamma = 10^{-2}\,h^{-1}$. b) Phage population density ($P$), bacterial population density ($B$), colony density $\rho$, and average colony size $\langle s \rangle$ as a function of bacterium dispersal rate $\gamma$ for a system of $V = 10$ ml for $\nu = 1/3$. c) Colony size distribution in steady state for $\gamma = 10^{-2}\,h^{-1}$. Simulation parameters are $\eta = 10^{-8}$ ml/h, $\lambda = 2\,\text{h}^{-1}$, $\delta = 0.1\,\text{h}^{-1}$, $\beta = 100$.

Based on the above observations, we can conclude that in our model, bacterial influx can be effectively substituted by the dispersal of bacteria from existing colonies. This provides a biologically relevant mechanism that better aligns with the dynamics of the natural microbial population.

### S4. Model dependence on value of exponent $\nu$

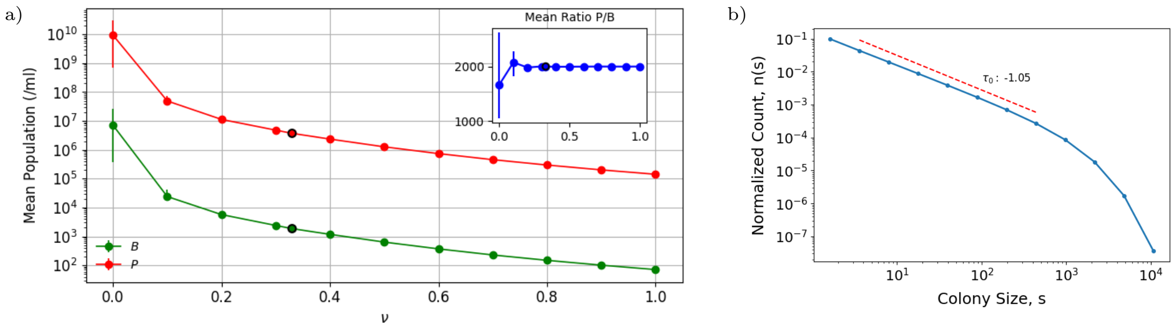

Figure S4: (a) Mean bacterial and phage populations as a function of $\nu$. The inset displays the phage-to-bacteria ratio, which remains constant. The highlighted data points correspond to $\nu = 1/3$, the value used in our model. Simulation parameters are $\alpha = 0.1\,\mathrm{ml}^{-1}\mathrm{h}^{-1}$, $\eta = 10^{-8}\,\mathrm{ml/h}$, $\lambda = 2\,\mathrm{h}^{-1}$, $\delta = 0.1\,\mathrm{h}^{-1}$, $\beta = 100$, and $V = 1\,\mathrm{L}$. (b) Colony size distribution for $\nu = 1$, showing a steeper exponential decay instead of the stretched exponential obtained for $\nu = 1/3$.

### S5. Spatial Heterogeneity

We consider a spatial grid of $L_x \times L_y \times L_z$ where each grid point is $dl = 10^3 \mu m$. Hence, each grid point has a carrying capacity of $10^9$ bacterial cells. Due to this factor, we consider limiting the growth of each microcolony in each grid cell. This assumption is reasonable considering that our microcolonies grow up to $10^6$ cells in the $\nu = 1/3$ case. We have also checked the model in main text for limiting growth, the behavior and $P/B$ ratio doesn't get affected by this assumption. We consider phages are diffusing in the 2d grid with diffusion constant D. Hence, the dynamics are governed by the following equations:

$$\begin{aligned} \frac{dB_i}{dt} &= \lambda B_i \left(1 - \frac{B_i}{K}\right) \quad for\ i \ = 1, 2, ...N \\ \frac{\partial P}{\partial t} &= D\left(\frac{\partial^2 P}{\partial x^2} + \frac{\partial^2 P}{\partial y^2}\right) - \delta P \end{aligned} \tag{S15}$$

For the low enough bacteria influx $\alpha$, the typical distance between colonies $d$ limits the time for phages locating a new colony by diffusion $\propto d^2/D$. In 2-dimension, $d^2 \propto 1/\alpha$, hence The $P/B$ ratio should be proportional to $1/\alpha$ for low $\alpha$. With the current parameter and $D \sim 5 \times 10^{-12}\,\mathrm{m}^2/\mathrm{s}$ [1], we confirmed that the $P/B$ ratio is proportional to $\alpha$ for $\alpha \lesssim \frac{10}{\mathrm{ml \cdot h}}$ and reached $\sim 10$ for $\alpha \sim \frac{0.1}{\mathrm{ml \cdot h}}$ (Supplementary Fig S5)

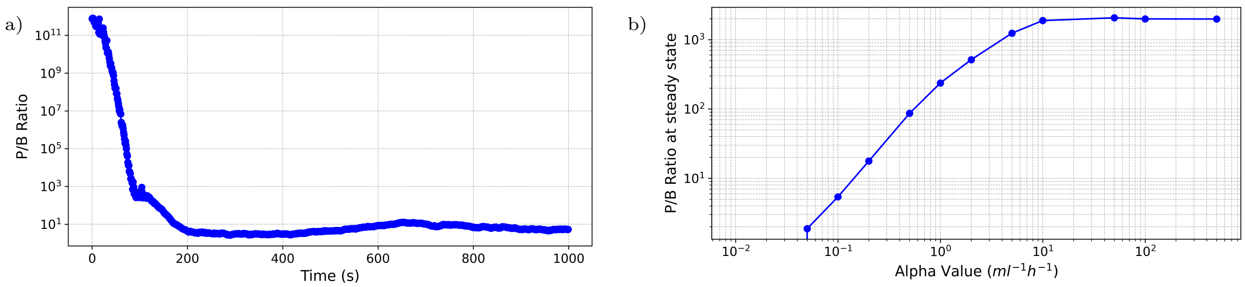

Figure S5: a) $P/B$ for spatially distributed colonies with limited growth ($\alpha = 0.1\,\mathrm{ml}^{-1}\mathrm{h}^{-1}$, $V = 1\,\mathrm{L}$). b) $\alpha$ vs $P/B$ .

As we increase $\alpha$, the system approximates to a well-mixed model. In this scenario, colonies are well distributed, and the diffusion of phages becomes insignificant in terms of overall dynamics.

### S6. Model with exponential decay of colonies

In the model in the main text, we consider the instantaneous burst of the colony as soon as it gets attacked. Here, we compare it with a model in which we consider the time delay between a colony being attacked and decaying. We compare two cases: a) instantaneous decay, b) colony decays exponentially with rate $\ln(10)h^{-1}$, a colony of size $10^n$ takes $n$ hours to decay. As the colony get attacked, it starts decaying/dying with a constant rate $\ln(10)h^{-1}$, following $dB_i/dt = -\ln(10)B_i$. So, at each time step,

$$B_i(t + \Delta t) = B_i(t)e^{-\ln(10)\Delta t}$$

the colony size is reduced by a factor of $e^{-\ln(10)\Delta t}$. The increase in phage population is given by,

$$P(t + \Delta t) = P(t) + \beta \sum (B_i(t + \Delta t) - B_i(t))$$

When $B_i(t)$ reaches less than size 1, the colony $i$ is removed from the system.

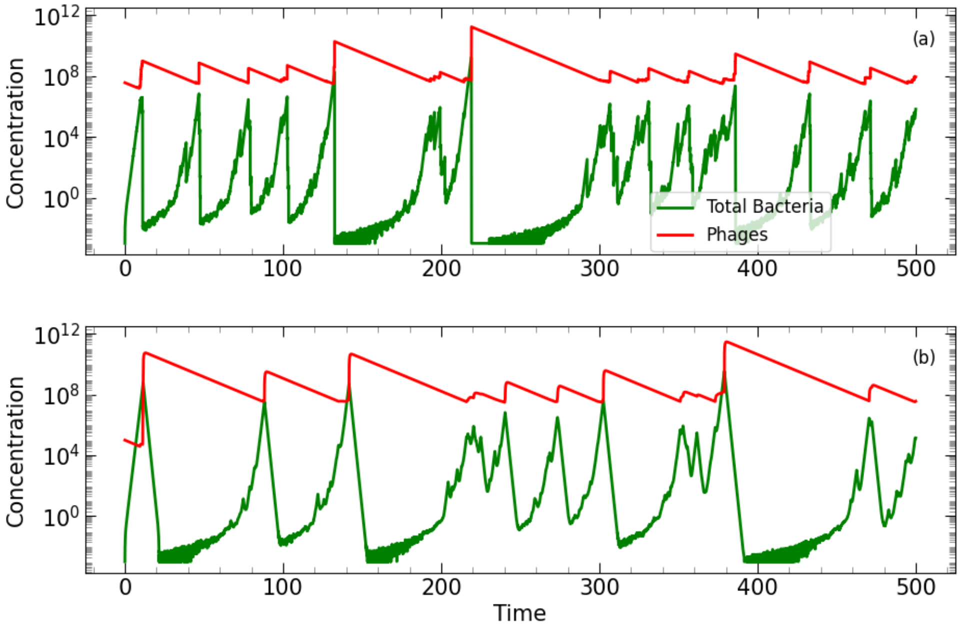

Figure S6: Temporal evolution of population for a) instantaneous burst of colony releasing phages in system b) Colony decays exponentially at rate $\ln(10)$ for $\nu = 0$.

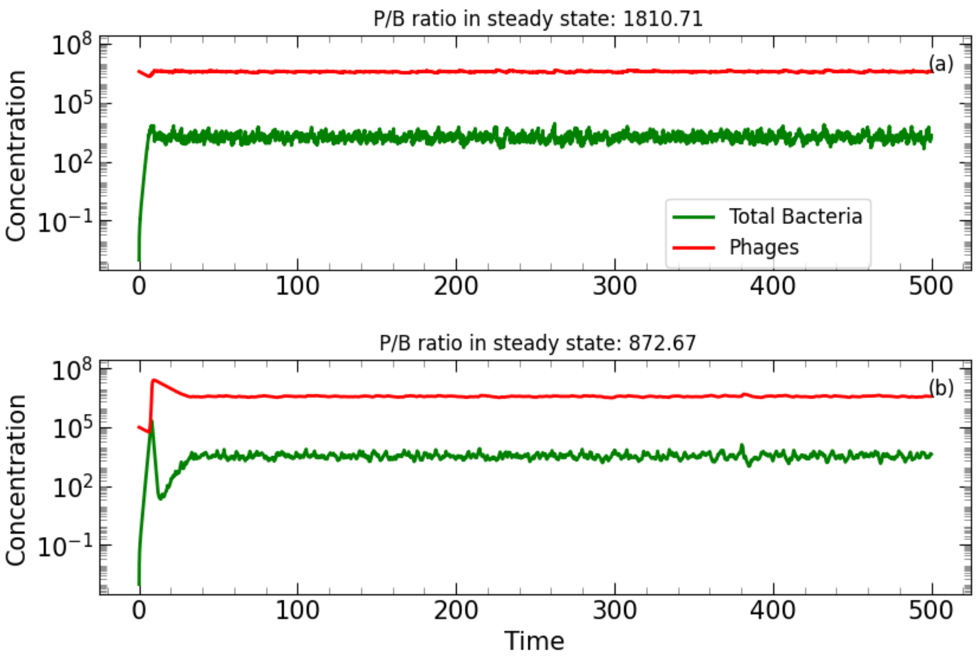

Figure S7: Temporal evolution of population for a) instantaneous burst of colony releasing phages in system b) Colony decays exponentially at a constant rate ln(10), for $\nu = 1/3$ volume, $V = 1L$, with simulation parameters $\alpha = 0.1h^{-1}ml^{-1}, \eta = 10^{-8}, \lambda = 2h^{-1}, \delta = 0.1h^{-1}, \beta = 100$

### S7. Fluctuations in $B$, $P$ and colony sizes with adsorption rate, $\eta$ and influx rate, $\alpha$

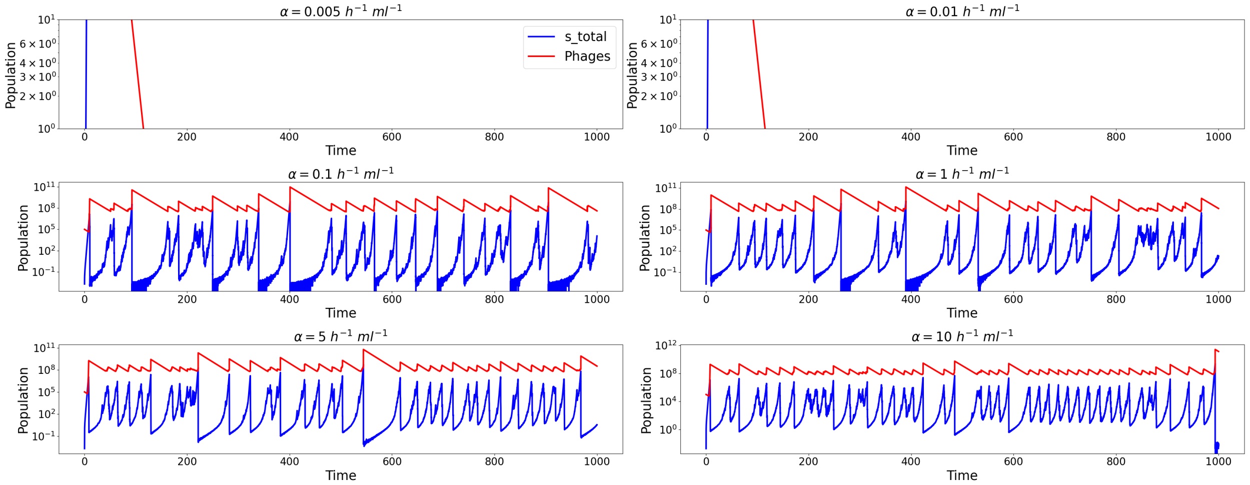

Figure S8: Temporal evolution of bacterial density ($B$) and phage density ($P$) for different values of the bacterial influx rate $\alpha$, under the condition $\nu = 0$ (infection rate independent of colony size). Higher $\alpha$ values stabilize the dynamics, as the increased influx balances phage predation, resulting in decreased amplitude of fluctuations in both bacterial and phage populations. In contrast, lower $\alpha$ leads to pronounced episodic fluctuations, with sharp increases and collapses of bacterial density driven by phage bursts.

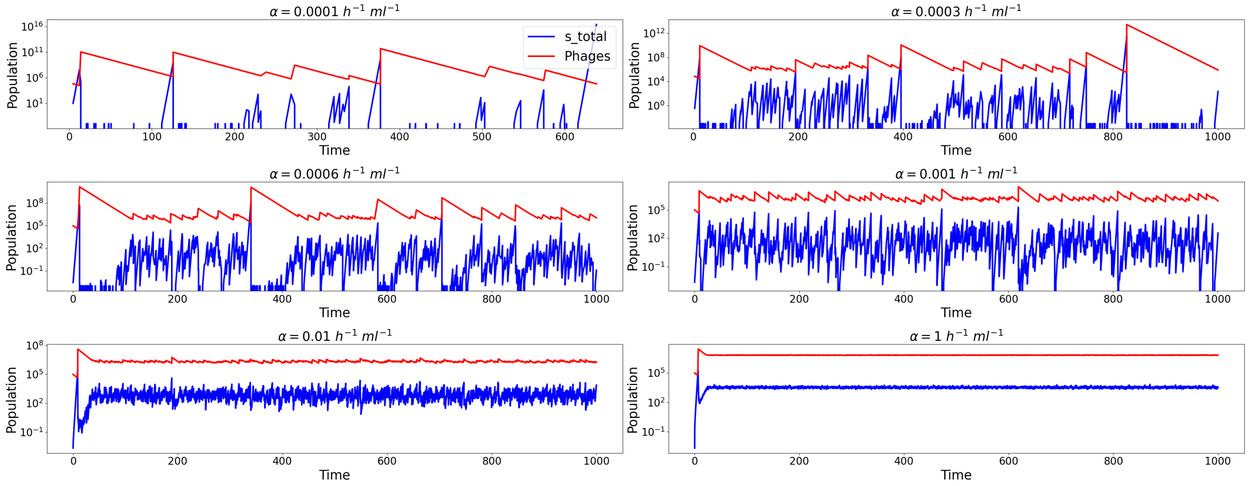

Figure S9: Temporal evolution of bacterial density ($B$) and phage density ($P$) for different values of the bacterial influx rate $\alpha$, with $\nu = 1/3$ (diffusion-limited infection rate scaling with colony size). The parameters are the same as those in the main text, with a system volume of $V = 1\,\mathrm{L}$. The inclusion of ($\nu = 1/3$) allows system to coexist for even lower $alpha$ compared to the $\nu = 0$ case (Fig. S8), where episodic dynamics dominate. Higher $\alpha$ reduces these fluctuations, stabilizing both bacterial and phage densities. This highlights how bacterial influx and $\nu$ jointly influence the stability and variability of population dynamics.

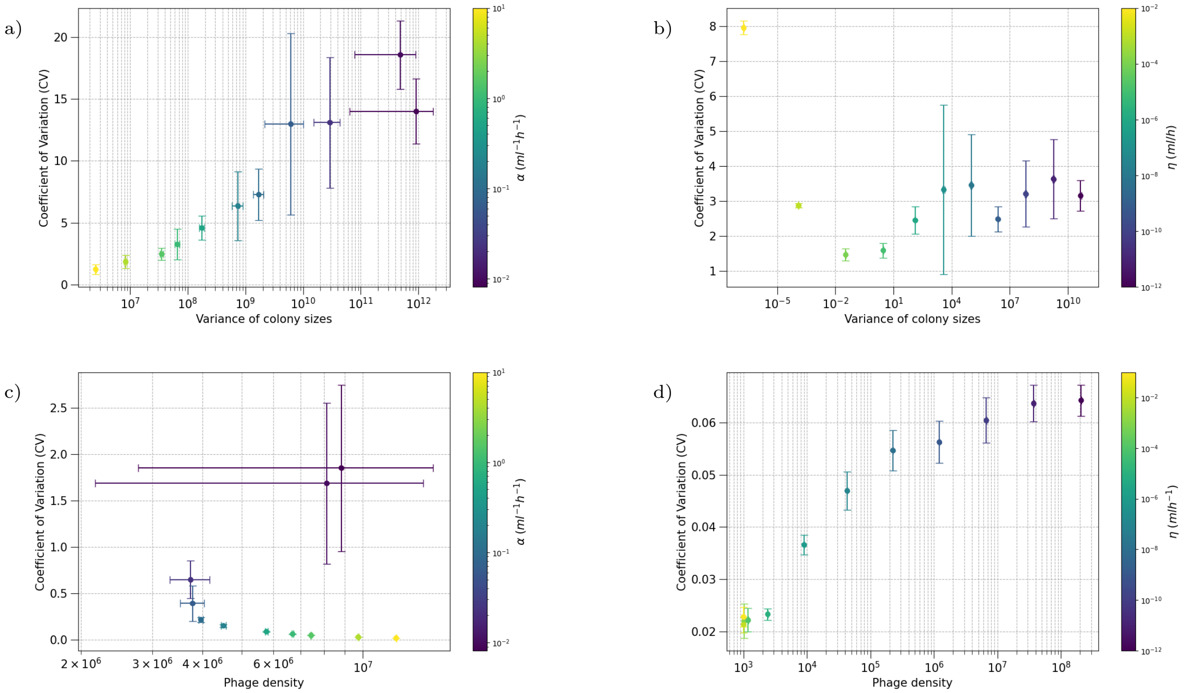

Figure S10: Fluctuations in the coefficient of variation (CV) for (a, b) variance of colony sizes and (c, d) phage density as a function of the influx rate $\alpha$ and the infection rate $\eta$ for $\nu = 1/3$. Panels (a) and (b) show that the CV increases significantly with higher variance in colony sizes, particularly at lower $\alpha$ and $\eta$, indicating pronounced stochastic effects in these regimes, as can be seen in the temporal evolution of phage and bacteria density in Fig. S9. Similarly, panels (c) and (d) illustrate the variability in phage density, where lower $\alpha$ and $\eta$ result in heightened fluctuations due to reduced system stabilization. Notably, higher $\alpha$ and $\eta$ suppress fluctuations, leading to more stable colony and phage population dynamics. This highlights the critical role of both parameters in regulating ecological stability and population variance, with smaller parameter values exacerbating noise-driven dynamics.
